# Urban colonization of invasive species on islands: *Mus musculus* and *Rattus rattus* genetics of establishment on Cozumel island

**DOI:** 10.1101/2023.07.14.549045

**Authors:** Gabriela Borja-Martínez, Ella Vázquez-Domínguez

## Abstract

Humans and wildlife experience complex interactions in urban ecosystems, favoring the presence of commensal species, among which invasive species are particularly successful. Rodents are the main vertebrate group introduced to oceanic islands, where the invasion process and dispersal patterns strongly influence their evolutionary and genetic patterns. We evaluated the house mouse *Mus musculus* and the black rat *Rattus rattus* on Cozumel island, Mexico. We assessed genetic diversity and structure, connectivity, gene flow, relatedness and bottleneck signals based on microsatellite loci. Our findings show that the constant introduction of individuals of different origins to the island promotes high allelic diversity and the effective establishment of migrants. We identified a clear genetic structure and low connectivity for the two species, tightly linked with anthropogenic and urban features. Moreover, we found *M. musculus* has a particularly restricted distribution within the city of San Miguel Cozumel, whilst its genetic structure is associated with the historical human population growth pulses accompanying the urbanization of the city. At the fine-scale genetic level, the main urban drivers of connectivity of the house mouse were both the impervious land surfaces, i.e. the urban landscape, and the informal commerce across the city (a proxy of resources availability). Chances of a secondary invasion to natural environments have been relatively low, which is crucial for the endemic taxa of the island. Nonetheless, improving urban planning to regulate future expansions of San Miguel Cozumel is of the outmost importance in order to prevent these invasive species to disperse further.

## Introduction

Cities worldwide vary in age, shape, size, human population, geographic context and history, and are undergoing accelerated expansion, making them highly dynamic and unstable systems. Urban ecosystems have high heterogeneity, usually characterized by mosaics of land use, including infrastructure with different purposes (residential, commercial, industrial), urban green space (parks, gardens) and natural remnants [1,2], frequently organized along gradients from the city centers to suburban surrounding landscapes. In these environments, humans and wildlife experience complex interactions, affecting the evolutionary trajectory of species [3-6]. Urbanization favors in particular the presence of closely human-related species, organisms that have changed life history features like breeding cycles, diet, behavior, or home ranges, enabling them to thrive in urbanized environments, and among which invasive species are especially successful [2,4].

Invasive (non-native) species are those that have been accidentally or intentionally transported and released outside of their natural range, have dispersed from the area of introduction and have established reproductive active populations [2-7]. In recent decades, global commercial exchange and transportation have facilitated and enhanced biological invasions. The invasion dynamics, i.e. propagule acquisition, introduction, establishment, proliferation and spread, greatly determines the abundance, structure and impact of invasive species [8,9]. Also, the introduction and dispersal patterns of non-native species strongly influence their genetic diversity and structure, demography, and natural selection [10-12]. Indeed, the success of invasion is limited when the founding populations are small, due to decreased genetic variability, genetic drift and inbreeding resulting from founder effects. In this scenario, the ‘new population’ becomes highly differentiated from the source population, potentially leading also to fixation of deleterious alleles [8,13]. However, genetic erosion can be avoided by the recurrent introduction of new alleles through multiple invasion events; also, if the number of initial propagules is high and the bottleneck is short, allows for a rapid subsequent population expansion and increased genetic diversity. Hence, the propagule pressure (i.e. relation between the propagule size and propagule frequency) is a key factor for the success or failure of an invasion [10,12,14,15]. The availability of resources in cities facilitates the establishment and proliferation of invasive species, while road networks allow the rapid dissemination of propagules [2,16]. In addition, multiple social, economic and ecological factors of cities influence their distribution and abundance, with varying effects on population genetic patterns, gene flow and connectivity of animals and plants [1,2,6,17]. Nonetheless, our understanding of the dynamics of invasion in urban ecosystems is still limited, particularly on islands.

Among invasive species, rodents are among the most widely introduced vertebrates. Their success is associated with their high reproductive rates, short generation times and dietary plasticity; they can consume organic matter, seeds, fruits, different invertebrate and small vertebrate species, eggs and coastal resources [18-23]. Additionally, their evolutionary history is tightly linked with their commensal ecology [1,24,25], to such a degree that they are considered the main group introduced to oceanic islands [26]. Moreover, their global dispersal is significantly associated with human colonization, expansion and urbanization, also regulated by the intensity of commercial and cultural ties established between human populations throughout their history [27]. This is the case of the Norwegian rat (*Rattus norvegicus*), the black rat (*Rattus rattus*) and the house mouse (*Mus musculus*).

The black rat is considered the most damaging invasive rodent on islands [28]. It evolved in the Indian peninsula on southern Asia, spread across Europe via the Roman trade (400 BC-100 AC), and dispersed into America and the islands of the Indian and Atlantic oceans with the explorations of the 16^th^ and 17^th^ centuries [29,30]. While *Mus musculus* originated in the Iranian Plateau and is characterized by three evolutionary lineages that diverged 100,000 years ago: *M. musculus castaneus* in Asia, *M. musculus musculus* which colonized central and eastern Europe 3,000 years ago, and *M. musculus domesticus* that established in western Europe and arrived in America, Africa and Australia associated with European expeditions beginning in the 15^th^ century [31-33]. There are many genetic studies of *R. rattus* and *M. musculus* focused on diversification, phylogeographic patterns and invasion routes (e.g. [30,32,34-39]), both in continental regions [33,40-42] and in islands [43-46]. Nonetheless, few studies have investigated their genetic and ecological dynamics associated with urban ecosystems specifically on islands [47].

Our aim was to evaluate the evolutionary and genetic patterns of two island invasive rodents, *Mus musculus* and *Rattus rattus* on Cozumel island, Quintana Roo, Mexico, associated with the urbanization history of the island. We assessed genetic diversity and structure, connectivity, gene flow, relatedness and contemporary and ancestral bottleneck signals of the two rodent species using microsatellites loci. In addition, given the particularly restricted distribution we found of the house mouse within the city of San Miguel Cozumel, we evaluated the species’ colonization process and connectivity based on demography and a landscape genetics framework. To this end, we integrated environmental, anthropogenic and social variables to adequately represent the urban ecosystem. Cozumel island is one of the main tourist destinations in Mexico and the busiest tourist port in the Caribbean, thus we assumed a constant introduction of individuals of both invasive species via the island seaports and, accordingly, predicted that both species will exhibit moderate genetic diversity values and weak bottleneck signals. Additionally, San Miguel Cozumel preserves some remnants of natural vegetation and, despite its small size and relatively high income from the tourist industry, has high economic heterogeneity and social inequality. Hence, we expected the two rodents populations will exhibit spatial genetic structure and low connectivity associated with urban and socioeconomic features. Yet, we predicted a differential response to urbanization for each species owing to their distinct life history and social organization, with a greater genetic differentiation in *M. musculus*. We show that considering the genetic history of introduction and past demographic events is key for the understanding of invasion processes of commensal rodents on islands.

## Materials and methods

### Study site and sampling

Cozumel (ca. 486 km^2^) is an oceanic island of coralline origin that has never been connected to the mainland [48,49]. It is located 17.5 km off the Yucatán peninsula in the Caribbean Sea (20°16’18.2’’-20°35’32.8’’N; 86°43’23.3’’-87°01’31.1’’W), separated from the mainland by the Cozumel Channel (400 m deep). It is the Mexican island with the highest number of endemics (31 taxa) and nearly 75% of its native vegetation [50]; conservation efforts have rendered two terrestrial (Refugio Estatal de Flora y Fauna Isla Cozumel and Reserva Estatal Selvas y Humedales de Cozumel) and one marine (Parque Nacional Arrecifal de Cozumel) protected areas. Despite this, a growing human population and increased cruise ships tourism have accelerated urbanization, habitat loss and fragmentation [51]. Consequently, numerous exotic species have been, and continue to be, introduced to the island (e.g. domestic cats and dogs, black rat, house mouse, boa, margay), which have become feral or invasive [50,52,53].

Cozumel island can be divided based on human-influence zones as: i) the city of San Miguel Cozumel, home to 98% of the island’s inhabitants [54]; it includes the airport, hotel developments on the north and south of the city, and main residential and commercial infrastructure, ii) the sanitary landfill on the east, and iii) several scattered small settlements, e.g. several ranches and El Cedral town (Fig 1).

**Fig 1.**
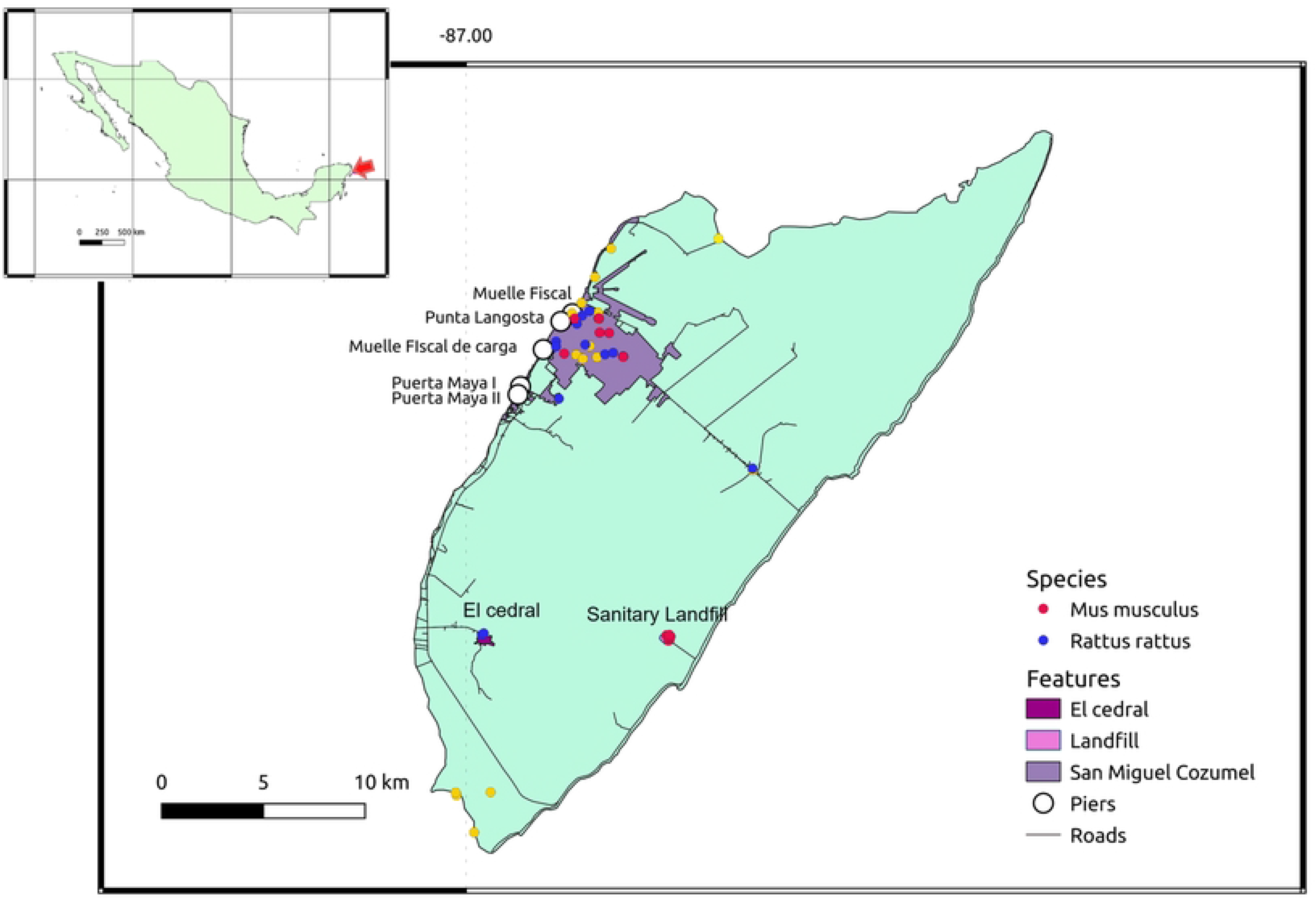
Map of the study site and sampling localities on Cozumel island. Sampling points are shown where *Mus musculus* (red dots) and *Rattus rattus* (blue dots) were sampled; yellow dots depict sites where no individuals were captured. The urban area of the main human settlement, San Miguel Cozumel city, is shown in pink. Other human-influence zones are indicated, the sanitary landfill and El Cedral town, the maritime terminal and the ferry port, and the Puerta Maya and Punta Langosta piers.

We sampled *Rattus rattus* and *Mus musculus* with Sherman-live traps during September 2015 in 12 neighborhoods within San Miguel Cozumel, plus the sanitary landfill, El Cedral town and a few small human settlements around the island. The number of traps per site and the number of houses sampled varied by neighborhood (S1 Table). All individuals were euthanized using an anesthetic chamber with ethyl ether, following the procedures allowed by the Mexican Official Norm [55]. Post-mortem ear tissue samples were immediately collected and preserved in Eppendorf tubes with 96% ethanol. We also collected tail bones of run over individuals.

### Microsatellite genotyping

We extracted genomic DNA from tissue samples with the Genomic DNA Tissue Miniprep kit (ZimoReserch), following the manufacturer’s instructions. For *R. rattus* we genotyped 10 nuclear loci, eight species-specific: Rr14, Rr17, Rr21, Rr22, Rr67, Rr68, Rr107, Rr114 [28] and two designed for *R. norvegicus*: D16Rat81, D5Rat83 [56]; we chose 10 specific microsatellites for *M. musculus*: D5Mit149, D6Mit309, D9Mit54, D13Mit61, D15Mit98, EGG22992, PP4A02, PP7B08, PP10A02, PP10E08 [35,46,57]. Microsatellites were amplified by PCR reactions (see protocol details in S2 Table). Fragment analysis was performed with the capillary sequencer AbiPrism3730xl Analyzer (Roy J. Carver Biotechnology Center, University of Illinois, USA) with the molecular weight marker Rox500 size standard. Fragment analysis and genotyping were done with GeneMarker v.2.6.7.

### Genetic structure, diversity, relatedness and bottlenecks

To infer genetic structure we used a Discriminant Analysis of Principal Components (DAPC) [58] with adegenet in R v.2.1.5 [59], a multivariate approach that reflects the genetic variation differences between groups while minimizing variation within them; we applied the alpha-score method to determine the proportion of successful reassignment by individual in function of the number of retained PCs (*R. rattus*: 1 PC; *M. Musculus*: 8 PCs). We also performed a spatial genetic analysis with GENELAND v.4.0.4 [60], a spatial Bayesian method based in Monte Carlo Markov chains and Poisson-Voronoi tessellations to determine genetic discontinuities; 30 independent runs with 10,000,000 interactions and 100,000 of burn-in were performed, an uncertainty coordinate value of 30 m for *R. rattus* based on its low dispersion in urban zones [20], while for *M. musculus* we used a 20 m value considering the species’ home range in cities [61].

The following analyses were performed for the entire population and for the different genetic clusters obtained for each species (see genetic structure Results). We screened for null alleles and stuttering in Micro-Checker v.2.2.3 [62], using 3,000 randomizations and adjusting the values for multiple comparisons with a Bonferroni sequential correction. We also tested genotypes for Hardy-Weinberg equilibrium and linkage disequilibrium with the Fisher exact test with 1,000 batches and 100,000 interactions in GENEPOP v.4.0 [63] and Bonferroni correction. Genetic diversity was estimated with GENALEX v.6.501 [64] as the average number of alleles per locus (*Na*), number of effective (*Ae*) and private alleles (*Ap*), observed (*Ho*), expected (*He*) and unbiased expected heterozygosity (*H_NEI_*) (corrected for small sample sizes; [65]), and inbreeding coefficient *F_IS_*. We assessed relatedness by estimating the likelihood of two individuals belonging to the same group being half-siblings, full siblings, parent-offspring or unrelated, with ML-RELATE [66].

To evaluate genetic bottlenecks we used two approaches, a graphical method to evaluate contemporary bottleneck, based on the distribution of allele frequencies in which recently bottlenecked populations are likely to have lost mainly the rare alleles [67]. We also tested for heterozygote excess with the program BOTTLENECK v.5.1.26 [68,69] for ancestral bottleneck, by comparing observed and expected heterozygosity values estimated under three mutational models: infinite alleles (IAM), stepwise mutation (SMM) and two-phase (TPM) with two variants (90% SMM, 10% IAM and 10% variance; 70% SMM, 30% and 10% variance). Models were run with 10,000 replicates and significance was calculated with a Wilcoxon test.

### Differentiation and migration

Pairwise genetic differentiation between genetic clusters for both species was estimated with *F_ST_*and *R_ST_* with ARLEQUIN v.3.1 [70]. Ancestral migration was assessed with an indirect estimator of gene flow (*Nm*), based on the allelic differences between genetic clusters (*F_ST_*), with the formula 4Nm = (1/*F_ST_*). The number of migrants was estimated both by individuals and by sexes with ARLEQUIN v.3.1. This model assumes a constant population size and symmetric migration rates [71]. We also estimated contemporary bidirectional migration rate using BAYESASS v.3.0. [71], with 5×10^6^ interactions, 3×10^5^ burn-in, and sample rate of 2,000. The delta value was adjusted with various trials for each species, based on which we chose values of 0.25 for *R. rattus* and 0.20 for *M. musculus*. Chain convergence was evaluated with the Faubet method [72], which calculates the Bayesian deviation based on the posterior probabilities that a certain genotype is possible under the ancestral migration rate and the posterior probability of allocation given the same migration rate [72,73].

### *Mus musculus* colonization patterns

For our objective to evaluate the colonization patterns and connectivity for *M. musculus* and explore the landscape genetics of the species, we specifically assessed if the urbanization growth pulses that occurred in San Miguel Cozumel determined the observed genetic structure and genetic diversity changes in the three genetic clusters obtained (SanMi1, SanMi2, SanMi3; see Results). First, we inferred the time, expressed as number of generations, and patterns of colonization with the Approximate Bayesian computation method (ABC) [74], implemented in DiyABC v2.04 [75]. We evaluated three simple scenarios built based on the history of human population presence on Cozumel, using datasets simulated by coalescent from prior distribution parameters [76]: i) Scenario1, we hypothesized a sequential colonization where SanMi1 is the ancestral (introduced) Cozumel population, from which SanMi2 originated and then SanMi3 from this one; ii) Scenario2, we tested if SanMi2 and SanMi3 originated either as two independent populations from SanMi1 or from two colonization pulses at different times; and iii) Scenario3, we evaluated if SanMi1 and SanMi3 originated independently from the same unsampled introduced population that colonized Cozumel, while SanMi2 originated from SanMi1 (Fig 2a). For all introduction and colonization events we assumed short genetic bottlenecks. *Mus musculu*s has a generational time of 2-3 months in captivity, but in the wild it can have only two litters per year, depending on resources availability [77,78].

**Fig 2.**
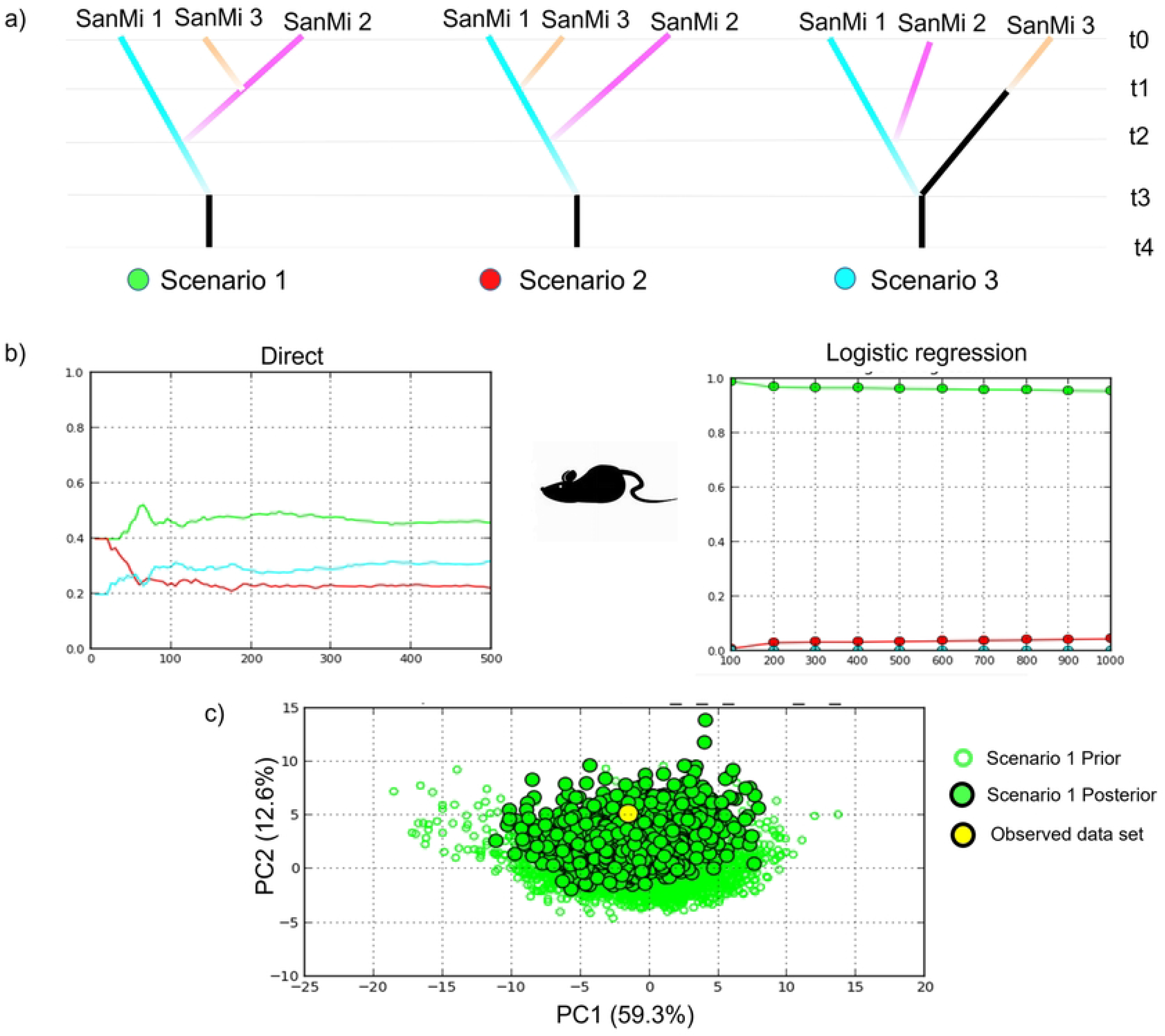
Colonization patterns of San Miguel Cozumel by *Mus musculus* based on the history of human population presence on Cozumel. **a**) scenarios tested with DiyABC v.2.04 [75]. Scenario 1: sequential colonization where SanMi1 is the ancestral (introduced) population-SanMi2-SanMi3; Scenario 2: SanMi2 and SanMi3 originated either as two independent populations from SanMi1 or from two colonization pulses at different times; Scenario3: SanMi1 and SanMi3 originated independently from the same unsampled introduced population, and SanMi2 originated from SanMi1. **b**) Scenario model comparison performance considering the direct and logistic approaches. **c**) Posterior predictive test of model fit corresponding to Scenario 1.

Originally inhabited by native Maya, Cozumel island was abandoned around 1570, undergoing a major population decline some decades after the Spanish conquest in 1521 [79]. More than three hundred years later, in 1847, people established again in Cozumel because of the indigenous uprising know as Caste War in the Yucatán peninsula, when rebellious Maya people escaped seeking refuge on the island [80]. We defined the priors of the divergence time between populations based on the elapsed times between the historical dates mentioned until 2015 and multiplied by the number of generations in one year. Hence, we defined priors accordingly for introduction of SanMi1, between 336 and 1008 generations. In 1980 Cozumel experienced a first population growth that promoted further urbanization, based on which the prior for SanMi2 was set between 70 and 210 generations; finally, another population growth wave started in 2000, so the prior for SanMi3 was between 30 and 90 generations (see below). We modeled the effective population size applying the default values [45,75] and applied the generalized stepwise mutation model for microsatellite data [81]. We generated 6×10^6^ simulated datasets per scenario, and scenarios were compared with the logistic and direct approaches using the following summary statistics: mean of number of alleles, genetic diversity and size variance within and between populations [12,81]. Confidence in the most supported scenario was evaluated by modeling 500 pods (pseudo-observed data) from the prior distribution to estimate the false allocation rate, where Type I error (i.e. probability in which the true scenario is rejected) is the proportion of pods for which the scenario does not have the highest posterior probability; and Type II error (i.e. probability of accepting a scenario when it is not true), is the proportion of pods of which the scenario with the highest posterior probability is not the scenario evaluated [81]. Based on the number of generations that coalesce supported by the best model (see Results), we estimated the dates of colonization using 4 and 6 years for the rodents generation time, considering that high food availability favours a high reproductive rate.

### Landscape genetics and connectivity

We generated environmental and anthropogenic variables at a resolution of 30×30 m. For the environmental variables, we used Landsat 8 images to calculate the normalized vegetation index (NDVI) in QGis 3.8, where low NDVI values indicate impervious land surface (i.e. man-made constructions, roads; [82]). For the anthropogenic variables we used the National Housing Inventory 2015 data [83] to generate three indices that measure marginalization, overcrowding and commercial businesses. The marginalization index was based on the number of essential services to which the population has access (concrete floors, running water, sewer system, sanitation services and street lighting); the overcrowding index represents the percentage of housing where more than three people share a room; and the commercial businesses denote the percentage of neighborhoods that have informal sector businesses in none, some, or all of their surrounding roads.

To obtain a measure of genetic differentiation for *M. musculus* across its distribution, we estimated the proportion of shared alleles with adegenet, an individual-based genetic distance that has no biological assumptions and can be used for inbreeding populations [84]. To generate resistance values for our environmental and anthropogenic surfaces we followed the optimization framework developed by Peterman [85], which uses maximum-likelihood population effects mixed models. We optimized each resistance surface with the function commuteDistance in gstudio v.1.5.2 in R [86], with three independent runs to verify parameters convergence. We determined the best-supported features by AICc (Akaike’s information criterion corrected for small/finite sample size; [87]). After running each surface individually, we applied a multivariate approach for which we built a set of hypotheses considering socioeconomic features most likely relevant for the distribution of the house mouse (Table 1).

**Table 1.** Composite hypotheses tested.

| Hypothesis | Surface |
| --- | --- |
| Economic development | NDVI + Commercial businesses |
| Poverty | Marginalization + overcrowding |
| Social Factors | Commercial businesses + Marginalization + overcrowding |
| Urbanization | NDVI + Commercial businesses + overcrowding |

Composite hypotheses of features that potentially affect connectivity of *Mus musculus* within the city of San Miguel Cozumel. NDVI: normalized vegetation index.

## Results

A total of 48 individuals of *R. rattus* (25 male and 17 females; six undetermined) and 58 of *M. musculus* (36 males and 17 females; five undetermined) were captured (S1 Table). The microsatellite locus D5Rat83 in *R. rattus* was monomorphic and was excluded from analyses. There was no evidence of linkage disequilibrium in *R. rattus* while in *M. musculus* it was only significant when the entire island was evaluated as a population (D5Mit149-D9Mit54 and D5Mit149-PP7B08) and for the genetic cluster SanMi2 (D13Mit61-EG22992). Five loci of *R. rattus* (Rr14, Rr17, Rr22, Rr68 y Rr107) and four of *M. musculus* (D5Mit149, D15Mit98, PP7B08, PP10E08) deviated significantly from Hardy-Weinberg equilibrium after Bonferroni correction. Only Rr14 in *R. rattus* and eight loci in *M. musculus* (except D6Mit309 and D13Mit61) showed evidence of null alleles, though no locus was shared between genetic clusters, thus all loci were considered for further analyses.

### Genetic patterns

Geneland results identified three genetic clusters (*K* = 3) for the black rat (Fig 3), which correspond to San Miguel Cozumel (Coz1), the sanitary landfill and El Cedral (Coz2), and the hotel development on the southwest (Coz3); in contrast, with the DAPC we found two distinctive groups, where Coz2 and Coz3 form a single cluster. All subsequent analyses for the black rat considered the three genetic clusters from GENELAND. For the house mice both GENELAND and DAPC identified three clusters, all distributed within San Miguel Cozumel, where SanMi1 is located on the northeast and includes the sanitary landfill, SanMi2 in the northwest, and SanMi3 in the south (Fig 4). Genetic diversity results showed a total of 56 alleles for *R. rattus* (Table 2), with an average of 6.5 alleles per locus and moderate heterozygosity (0.601). Coz3 was the cluster with less genetic diversity. Regarding *M. musculus*, diversity was markedly higher with a total of 100 alleles (10 per locus), where SanMi3 exhibited the lowest diversity values (Table 2). All clusters for both species showed positive *F_IS_* values indicating inbreeding. Notwithstanding, relatedness results showed a high percentage of unrelated individuals (83.4% and 86.7%, *R. rattus* and *M. musculus* respectively) (Table 2). The ancestral bottleneck test was not significant under any mutational model, while the contemporary bottleneck analysis showed rare alleles in low proportion for Coz3 and SanMi3, suggesting a bottleneck signal or a result of low sample size (S3 Table, S1 Fig).

**Fig 3.**
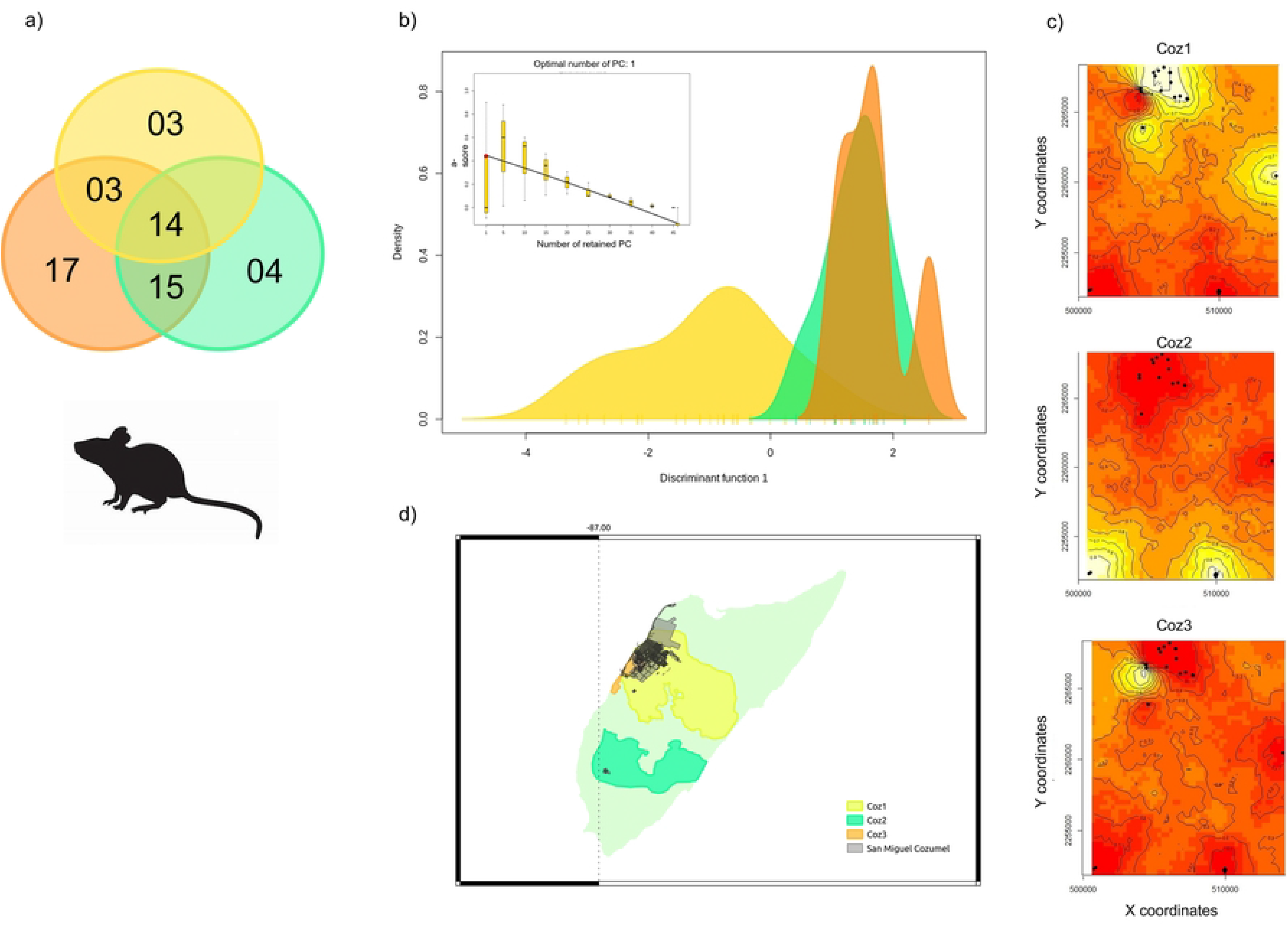
*Rattus rattus* genetic structure on Cozumel island. **a**) Venn diagram showing private and shared alleles between genetic clusters identified with GENELAND v.4.0.4 [60] (Coz1: yellow, Coz2: green, Coz3: orange). **b**) Scatter plot based on the GENELAND clusters as priors and the a-score plot used to determine the number of PCs (1 in this case) and discriminant functions to retain for the DAPC. **c**) Posterior probability maps belonging to each GENELAND genetic cluster where yellow colors depict the high probability isoclines. **d**) Map of the study site showing the distribution of the three genetic clusters considering the isocline = 0.5.

**Fig 4.**
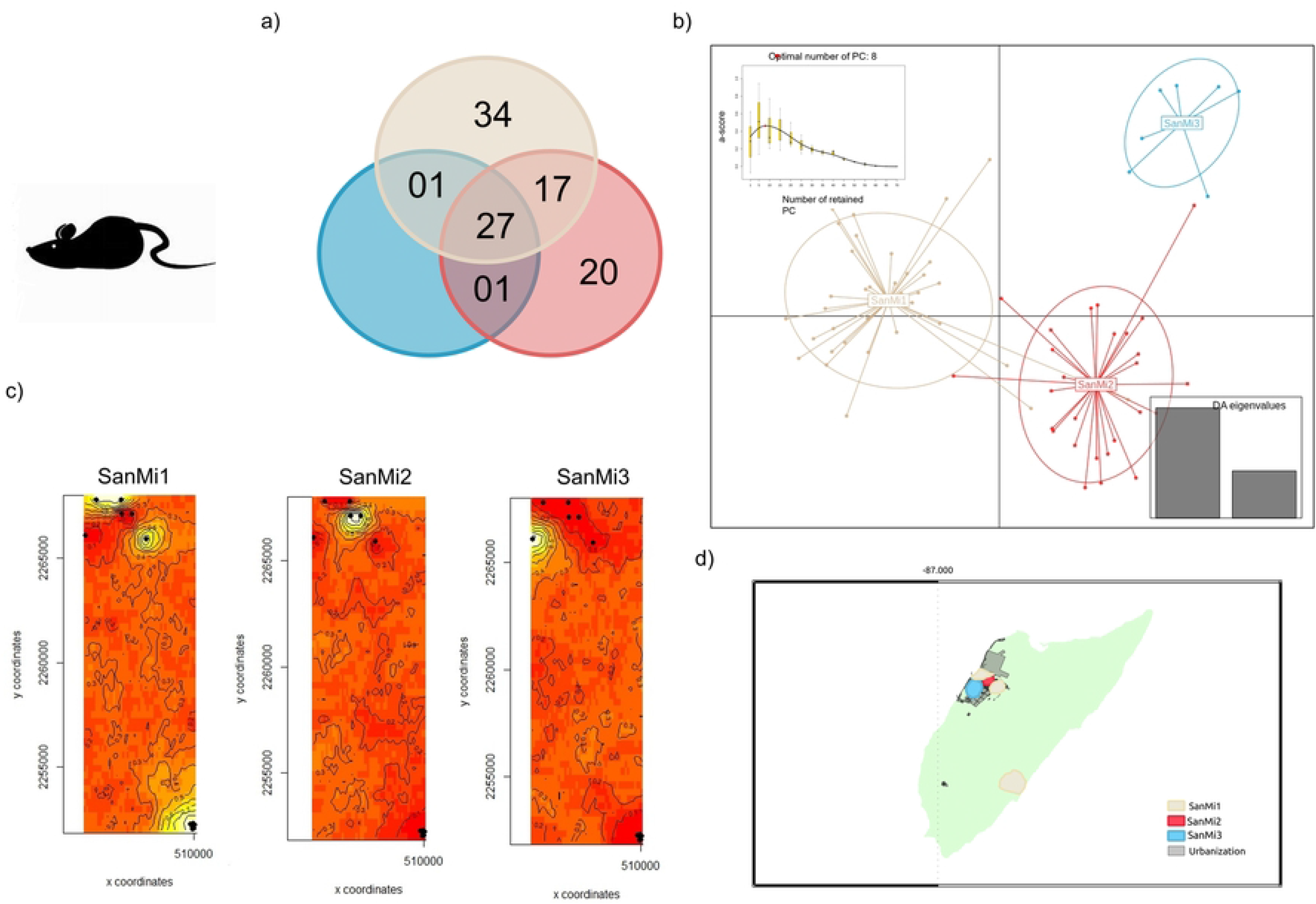
*Mus musculus* genetic structure on Cozumel island. **a**) Venn diagram showing private and shared alleles between genetic clusters identified with GENELAND v.4.0.4 [60] (SanMi1: brown, SanMi2: red, SanMi3: blue). **b**) Scatter plots based on the GENELAND clusters as priors and the a-score plot used to determine the number of PCs (8 in this case) and discriminant functions to retain for the DAPC. **c**) Posterior probability maps belonging to each GENELAND genetic cluster where yellow colors depict the high probability isoclines. **d**) Map of the study site showing the distribution of the three genetic clusters considering the isocline = 0.5.

**Table 2.** Genetic diversity and relatedness results for *R. rattus* and *M. musculus*.

| Genetic cluster | N | Na | Ae | Ap | Ho | He | $H_{NEI}$ | $F_{IS}$ | PO | FS | HS | NR |
| --- | --- | --- | --- | --- | --- | --- | --- | --- | --- | --- | --- | --- |
| *Rattus rattus* |  |  |  |  |  |  |  |  |  |  |  |  |
| Coz 1 | 28 | 5.889 | 3.138 | 17 | 0.494 | 0.581 | 0.591 | 0.140 | 0.013 | 0.024 | 0.011 | 0.85 |
| Coz 2 | 15 | 3.667 | 2.199 | 4 | 0.430 | 0.472 | 0.488 | 0.116 | 0.038 | 0.028 | 0.095 | 0.83 |
| Coz 3 | 5 | 2.222 | 1.784 | 3 | 0.289 | 0.371 | 0.412 | 0.233 | 0.0 | 0.10 | 0 | 0.90 |
| Cozumel | 48 | 6.556 | 3.366 | - | 0.447 | 0.595 | 0.601 | 0.233 | 0.014 | 0.034 | 0.117 | 0.83 |
| *Mus musculus* |  |  |  |  |  |  |  |  |  |  |  |  |
| SanMi 1 | 29 | 7.900 | 4.138 | 34 | 0.536 | 0.732 | 0.745 | 0.259 | 0.002 | 0.030 | 0.094 | 0.87 |
| SanMi 2 | 22 | 6.600 | 3.713 | 20 | 0.659 | 0.714 | 0.731 | 0.071 | 0.012 | 0.038 | 0.095 | 0.85 |
| SanMi 3 | 7 | 6.900 | 1.919 | 0 | 0.421 | 0.448 | 0.483 | 0.027 | 0.047 | 0.047 | 0.047 | 0.86 |
| Cozumel | 58 | 10.00 | 4.621 | - | 0.569 | 0.768 | 0.775 | 0.255 | 0.003 | 0.034 | 0.094 | 0.86 |

Summary of genetic diversity and relatedness for *Rattus rattus* and *Mus musculus* by genetic clusters and global. Number of genotyped individuals (N), average number of alleles per locus ( $N_a$ ), number of effective ( $A_e$ ) and private alleles ( $A_p$ ), observed ( $H_o$ ), expected ( $H_e$ ) and Nei ( $H_{NEI}$ ) heterozygosity, and inbreeding coefficient ( $F_{IS}$ ). Proportion of related individuals from the relatedness analyses (PO: parent-offspring; FS: full siblings, HS: half siblings and NR: no related).

### Ancestral and contemporary migration

Pairwise genetic differentiation between clusters was moderate to high (*R. rattus F_ST_* = 0.107-0.309, R*_ST_* = 0.086-0.52; *M. musculus F_ST_* = 0.094-0.184, R*_ST_* = 0.122-0.274 *p* < 0.05) (Fig 5a); the higher differentiation was observed between Coz2-Coz3 and SanMi1-SanMi3, respectively. Ancestral migration showed the highest exchange of individuals between Coz1-Coz2 (2.084) and SanMi1-SanMi2 (2.39) (Fig 5b). Distinct patterns were observed for each species regarding differential migration between sexes, where *R. rattus* females disperse more and, contrastingly, *M. musculus* males showed higher migration than females (Fig 5a). Whereas contemporary bidirectional migration results showed extremely low migration rates in *R. rattus* where at least 95% of individuals are permanent residents; the same was observed for *M. musculus*, where only the SanMi2 showed a comparatively lower permanent resident rate (0.68), while receiving a high proportion of migrants from SanMi3 (0.30) (Fig 5c).

**Fig 5.**
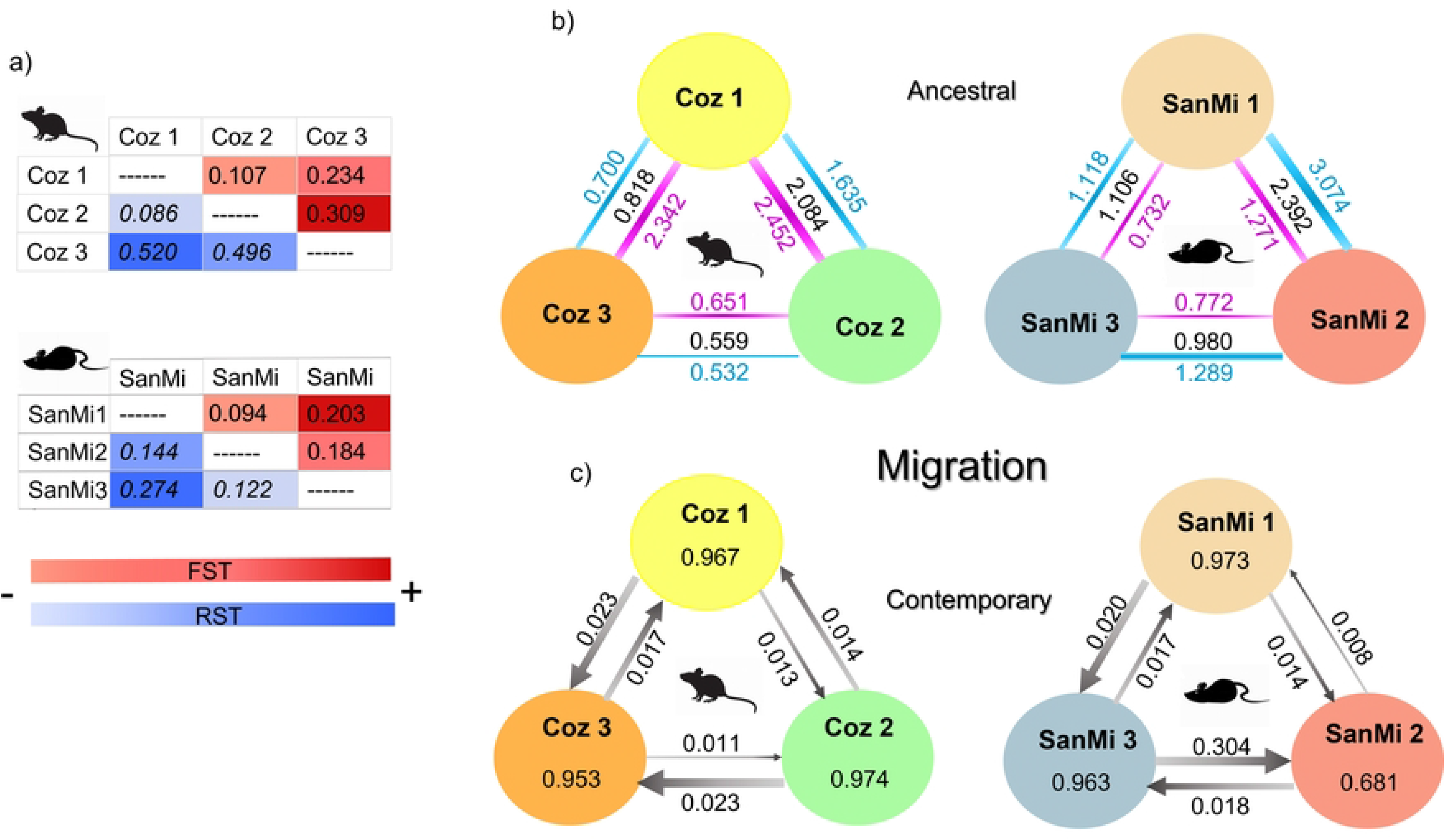
Differentiation and migration patterns of *Mus musculus* and *Rattus rattus* on Cozumel island. **a**) Pairwise genetic differentiation between genetic clusters based on *F_ST_* (red) and *R_ST_* (blue) estimators. **b**) Ancestral migration rates (*Nm*) based on the allelic differences between genetic clusters (*F_ST_*). The thickness of the lines and arrows varies depending on the intensity of migration between clusters. The color of the lines indicate the values for females (pink), males (blue), and global (black). **c**) Bidirectional contemporary migration rates estimated with BAYESASS v.3.0 [71], where the proportion of non-migrant individuals by generation per genetic cluster (within the ovals) and the direction of migrant individuals by generation between genetic clusters (next to each arrow) are shown.

### *Mus musculus* colonization patterns and landscape connectivity

Model comparisons resulting from the Approximate Bayesian computation showed that Scenario1, in which the sequential colonization was tested, had the highest probability based on both the direct (P=0.468, 95% confidence interval [CI] = 0.031-0.905) and the logistic (P=0.964, [CI] = 0.929-0.994) approaches (Fig 2b). The confidence test for the best scenario showed that type 1 error was 0.134 for the logistic approach. We performed a pre-evaluation of simulated scenarios via PCA and results indicated that the observed data fall within the cluster of simulated data points. In addition, the goodness-of-fit test using the posterior predictions exhibited that summary statistics from the simulated datasets based on Scenario 1 closely matched the empirical data (Fig 2c). Hence, based on the best model results and the generation times (4-6 reproductive events by year), we estimated the date of colonization events; accordingly, SanMi1 arrived to the island 676 generations ago (1846-1902), the divergence between SanMi1 and SanMi2 occurred 247 generations ago (1954-1974), and SanMi3 was founded 93 generations ago (1991-1999).

Results of model selection of the optimization resistance surfaces (environmental and anthropogenic) showed that the best-supported univariate model was geographic distance, identified as the top model 99% of the time, followed by NDVI (Table 3). The optimized NDVI surface assigned high resistance at values above zero that correspond to vegetated areas, and low resistance to bare landscape or urbanizations (S2 Fig). The linear mixed-effects models test of the composite resistance hypotheses showed that the best-supported multivariate model was Development (100% of the time; Table 4), where NDVI contributed the most to the model (88%).

**Table 3.** Univariate optimized model selection for *Mus musculus*.

| Surface | K | Equation | AICc | Average weight | Average rank | Top model (%) |
| --- | --- | --- | --- | --- | --- | --- |
| Distance | 2 | ----- | -771.65 | 0.896 | 1.00 | 99.53 |
| NDVI | 4 | Inverse Ricker | -766.82 | 0.096 | 2.10 | 0.25 |
| Commercial businesses | 6 | ----- | -759.04 | 0.006 | 3.04 | 0.21 |
| Overcrowding | 8 | ----- | -752.88 | 0.000 | 4.26 | 0.00 |
| Marginalization | 8 | ----- | -752.34 | 0.000 | 4.58 | 0.00 |

Univariate optimized model selection results based on maximum-likelihood population effects mixed models, for *Mus musculus* from the city San Miguel Cozumel. K: number of parameters used for the transformation of continuous surfaces plus the intercept; AICc: Akaike's information criterion corrected for sample size.

**Table 4.** Multivariate surfaces optimized for *Mus musculus*.

| Hypothesis | K | AICc | TopM (%) | Surface | Cmodel (%) |
| --- | --- | --- | --- | --- | --- |
| Development | 9 | -751.189 | 100 | NDVI | 88.16 |
|  |  |  |  | Commercial establishments | 11.83 |
| Poverty | 15 | -712.092 | 0 | Overcrowding | 50.71 |
|  |  |  |  | Marginalization | 49.28 |
| Urbanization | 16 | -704.088 | 0 | NDVI | 88.53 |
|  |  |  |  | Commercial establishments | 7.19 |
|  |  |  |  | Overcrowding | 4.27 |
| Social Factors | 20 | -642.977 | 0 | Commercial establishments | 31.57 |
|  |  |  |  | Marginalization | 33.53 |
|  |  |  |  | Overcrowding | 34.88 |

*Mus musculus* model selection results for the multivariate surfaces optimized. The contribution percentage (CModel) of each variable to the hypothesis and the resulting top model (TopM) based on bootstrap interactions are indicated. K: number of parameters used for the transformation of continuous surface plus the intercept; AICc: Akaike's information criterion corrected for sample size.

## Discussion

### Genetic diversity, invasion models and the genetic paradox of biological invasion

As a result of the reduced number of individuals with which introduced populations are established (founder effect), populations are expected to exhibit low genetic diversity, an effect that is theoretically stronger on islands [88,89]. However, in most invasive species this does not happen, which is known as the genetic paradox of biological invasion [8,9,89]. In agreement with this paradox, *R. rattus* showed moderate genetic diversity levels (*Ho*=0.44, *He*=0.59, *Na*=6.5), with higher levels in *M. musculus* (*Ho*=0.56, *He*=0.76, *Na*=10). These values are within the range found in other islands, like the black rat in Guadeloupe (*He*=0.69, *Na*=5), Madagascar (*He*=0.72) and New Zealand (*He*=0.75, *Na*=5.7) [43-45], as well as the house mouse on islands of New Zealand (*He*=0.51-0.56), Kerguelend (*He*=0.40, *Na*=2.8), Madagascar (*He*=0.67, *Na*=9.7) and Cyprus (*He*=0.77, *Na*=9.7) [26,34,35], despite the comparatively small size of Cozumel island. Interestingly, continental populations, some within cities, show similar diversity values [11,27,90].

Such levels of genetic diversity can be associated with the historical and demographic processes experienced by both the source and founder populations [91], namely the likely short ancestral bottleneck supported by our analyses, followed by rapid population growth, and coupled with multiple introduction events facilitated by the constant arrival of commercial and, more recently, cruise ships. In fact, a high number of private alleles as observed in the Cozumel populations may be the product of constant introductions of individuals of different origin, incorporating different genotypes into the genetic pool of the resident population and increasing allelic diversity [8,91]. This incorporation of new genotypes is favored by the ‘invasive meltdown model’, which establishes that ecosystems are more easily invaded as they are modified by previously arrived invasives, increasing population density [92-94]. In addition, the ‘invasive bridgehead effect’ refers to secondary invasions stemming from a particularly successful invasive population, which serves as a source of colonists for potentially new territories, increasing their probability of successful establishment [8,91]. These scenarios apply for most human settlements, as is the case in Cozumel, where the constant introduction of individuals of different origins into the island entails a strong propagule pressure, rendering high allelic diversity and the effective establishment of migrants. Similar examples include the house mouse in the Azores islands, which shows high allelic diversity resulting from multiple origins identified from southern Spain, Portugal, Finland and the Canary islands [38].

Independent colonization events from different lineages can be another factor influencing allelic differentiation and structure, as shown by *M. musculus* in the Kerguelen archipelago and the Canary islands, where the observed genetic diversity derives from the invasion of two distinct lineages [35,37], and the black rat in Madagascar that comprises two genetic groups product of two independent introduction events from the same ancestral population [45]. Our results do not support an independent colonization origin for the Cozumel clusters; instead, more recent social and demographic processes may also be contributing to the currently observed genetic diversity and structure.

### Genetic structure, urbanization and human population growth

As we predicted, the two rodent species exhibit a clear genetic structure. In our fine-scale genetic study, the three clusters identified for *R. rattus* correspond with three areas of the island with different anthropogenic impact (Fig 3), Coz1 comprises San Miguel Cozumel, Coz2 on the south of the island includes the sanitary landfill and the small El Cedral town; and Coz3 is located on the southwestern hotel zone, adjacent to the city, characterized by a combination of highly developed infrastructure surrounded by partially preserved forest remnants. These structure patterns contrast with what has been observed in other studies, for instance *R. rattus* did not show genetic differentiation in the Puketi Forest, New Zealand despite the marked habitat differentiation and significant geographic distances between samples sites [44], neither in Franceville, Gabon, a small African tropical city that has undergone a recent growth, regardless of the composite landscape including urban forest remnants and savannah [90].

Three genetic groups were also identified for *M. musculus,* but unlike *R. rattus*, these are tightly localized within San Miguel Cozumel, although SanMi1 also includes the sanitary landfill (Fig 4). These genetic clusters are limited both by roads and high traffic avenues and by the margins of the urbanized area. There are numerous examples in which avenues or highways constitute important barriers, for example in Salvador, Brazil where gene flow of the Norway rat (*Rattus norvegicus*) is limited by a main avenue [95]. Furthermore, *M. musculus* genetic structure is likely associated with the human population growth pulses accompanying the urbanization of San Miguel Cozumel. The spatial distribution of SanMi1 coincides with the limits of the city before 1970, when the tourism development began and the first large-scale contemporary human migration took place. Also, the site where the sanitary landfill is currently located was initially an open-air dump, thus the constant transport of waste trucks from the city to the dump prior to 1970 [51] can explain why the SanMi1 cluster extends to this area. As the urban sprawl grew, the house mouse was able to disperse and to establish new populations, so that SanMi2 includes all the new neighborhoods built between 1970 and 2000. Last, SanMi3 is distributed in the neighborhoods most recently established since 2000 onwards. Outstandingly, this relationship between the species genetic structure and urban growth coincides with the ancestral migration rate; that is, gene flow between the oldest clusters (SanMi1 and SanMi2) is greater compared to that between them and the most recent cluster (SanMi3). This sequential pattern also matches the genetic diversity of each cluster. Indeed, the ABC analysis supported our Scenario1 hypothesis of sequential colonization, where SanMi1 is the ancestral Cozumel population, which harbors the highest genetic diversity; from which SanMi2 originated and, more recently, SanMi3 that has the lowest genetic diversity. Loss of genetic variability through sequential colonization routes due to successive bottlenecks is frequent, as evidenced by the *M. musculus* invasion in Senegal [33].

The demographic growth of Cozumel is divided in two phases, a stationary one since the recolonization until 1950, and the tourism expansion from 1950 onwards, characterized by a constant population growth due of the intensification of human migration processes [96,97], which resulted in a disorganized urbanization expansion. In agreement with the spatial location of the genetic clusters, our best demographic model indicated that the initial colonization of Cozumel island occurred in SanMi1 between 1846 and 1902. In 1846, the Mayan uprising started in the Yucatan peninsula when Mayan farming refugees from the east of the peninsula moved to Cozumel; they founded two towns, San Miguel and El Cedral. The arrival of Mayan people to the island continued until 1913, thus these historical processes could be linked to the initial introduction of both invasive rodents. The origins of SanMi2 occurred between 1954 and 1974 coinciding with the initial urbanization pulse driven by tourism. Diving and fishing tourism were promoted in 1950 but is not until 1970 that a trust is created to promote tourist and economic development, initiating the construction of the northern hotel zone coupled with a strong demographic growth [80]. Finally, the colonization of SanMi3 occurred between 1991-1999, when Cozumel joined the Caribbean cruise network [51].

With this in mind and considering that Cozumel is the main port of the Mexican Caribbean, the high number of exclusive alleles in the different genetic clusters is highly relevant. Coz1 had the highest number of private alleles (17) for *R. rattus*, and SanMi1 (34) and SanMi2 (20) for *M. musculus*. A note of caution is that the low sample sizes for Coz3 and SanMi3 might underestimate the number of private alleles. Nonetheless, the clusters with highest number of private alleles occur in strategic areas of commercial exchange and tourism, supporting the hypothesis of a constant introduction of individuals to Cozumel. SanMi1, located in the oldest city square, includes also the maritime terminal (Muelle fiscal) and the cargo ferry port, the main passenger and commercial connection between the mainland and the island until the 1980s, when it started to receive cruises and two grand cruise piers were built, Puerta Maya (1997) and Punta Langosta (2000) [98], the latter also within SanMi1. Puerta Maya is adjacent to the southwestern hotel zone where SanMi3 and Coz3 are situated (Fig 1). Further phylogeographic and genetic assessments are needed to more finely reconstruct the routes of invasion of these rodents into Cozumel, but we hypothesize that Coz1 and SanMi1 share a genetic ancestry related to mainland individuals, particularly from Playa del Carmen that has historically been the region with the highest interchange with the island. More recent introductions would certainly include lineages from the various Caribbean ports that constitute the main Caribbean cruise route, i.e. Belize, Roatán (Honduras), Limón (Costa Rica), Colón (Panama), Santa María (Curaçao), Montego, Falmouth, Ocho Rios (Jamaica), and Grand Cayman ports [99].

The lack of ancestral and contemporary bottleneck signals in most of the genetic clusters for both species suggests a brief bottleneck after the introduction, a common pattern in species with high reproductive rate, short generation time and strong propagule pressure. In addition, the continuous introduction of individuals, from the same or different geographic origin, contributes to erasing the initial bottleneck signal and increases the global genetic diversity [12]. The house mouse has the ability to establish rapidly, as evidenced by the experimental introduction of two founders (one female and one male) on Saddle Island, New Zealand where the system’s load capacity was reached in only six months [100]. Likewise, the Norway rat has the ability to rebound population sizes after strong bottlenecks [101]. Yet, contemporary bottleneck signals were detected in SanMi3 and Coz3. As mentioned, the colonization of SanMi3 is relatively recent, thus likely still echoing the bottleneck signal, whereas such signal in Coz3 (located along the southwestern hotel zone) can be associated with the permanent pest control carried out by hotels, causing population size reductions and decreasing genetic variation. Indeed, significant loss of genetic variation related to control strategies has been documented in Poirier island and in Salvador, Brazil, as a result of a failed eradication plan for *R. rattus* [102] and after urban rat control campaigns [101], respectively. However, this would require further assessment because the SanMi3 and Coz3 bottleneck signals could also be associated with their low sample size.

### Ancestral and contemporary migration patterns

The dispersal capacity of a species depends, among others, on body size, morphology and physiology, as well as on biotic and abiotic factors like resources availability, habitat quality and geographic barriers. Dispersal can also vary between sexes as seen in birds and mammals, where the differential response of males and females relies on population density, competitive ability and habitat quality, among others, as a strategy to avoid inbreeding [103].

*Rattus* species are considered mostly philopatric, i.e. tending to return to or remain near a particular site or area, observed in females as is more common [20] but also found in males [104]. In our study in Cozumel, the black rat showed differential ancestral migration with higher values in females, supported by the findings of [104] in *R. norvegicus* at global scale; this could be associated with their social structure, where males have a strong hierarchy and control territories through which the females ‘freely’ roam; in fact female dispersion is food-determined and shapes male dispersion [105]. Accordingly, in the past females could have dispersed to sites with high resources availability, in agreement with the high gene flow between Coz1 and Coz2. However, contemporary gene flow showed extremely low migration between clusters; if the system is saturated (e.g. low resources, high competition) that would prevent the arrival of new migrating individuals. Only Coz3 receives some migrants from both Coz1 and Coz2, which could be explained by the population control measures implemented by hotels in this area, temporarily reducing competition for resources and allowing new migrants to arrive.

*Mus musculus* individuals tend to be nomads if population densities are low, but at high densities males are territorial [4]. This behavior patterns might have facilitated the presence of the three genetic groups within San Miguel Cozumel, with overall low ancestral migration. Contrary to the black rat, migration in the house mouse was higher for males, concordant with their social system where dominant males control and monopolize feeding sites, shared with several females and some subordinate males [31,61]. However, because population densities are usually high, such social hierarchies do not last long and males migrate constantly. As a result, populations behave panmitically on a regional scale but are structured on a finer scale [106]. Notably, as females are philopatric and do not move freely between territories, mitochondrial DNA-based studies have revealed that populations can be resistant to secondary female invasions [38], also explaining the observed low migration rate in this species.

### Urban drivers of connectivity in the house mouse

Connectivity analyses allow identifying key landscape elements needed to maintain genetic diversity and evolutionary processes. In human-related-species they also help to elucidate dispersal and distribution patterns of invasive species [26,27]. The most significant environmental feature influencing *M. musculus* connectivity was geographic distance, while the next best supported model was vegetation (NDVI). High and continuous vegetation cover is a good predictor of connectivity in natural environments, for example in rodents like *Peromyscus melanotis* [107], and in cities with non-human associated species like *Peromyscus leucopus* [108-109] and the lizard *Podarcis muralis* [110].

Contrastingly, as we predicted for this human-associated species, we found high connectivity for *M. musculus* along impervious land surfaces, i.e. the urban landscape, indicated by the low NDVI values recovered by the optimization surfaces. Similarly, Stragier et al. [27] found low gene flow in this species associated with highly vegetated areas in Dakar. Additionally, Fusco et al. [111] highlight that the variables analyzed in urban landscapes under a landscape genetics approach generally include human population density, roadways, infrastructure, impervious surfaces and industrial development, while social, cultural or political factors are rarely used. Indeed, incorporating socioeconomic variables was key in our approach. The best-ranked model in the multivariate analysis was our Development hypothesis, which considers the interaction between NDVI and Commercial businesses, further supporting that *M. musculus* connectivity is tightly linked with urbanization. The information used to build the Commercial businesses layer included all the informal commerce across the city, most of which is dedicated to street food selling. Based on the fact that they produce abundant waste, we used this as a proxy of resources availability for the house mouse. Interestingly, Combs et al. [112] show that low food waste availability decreased the genetic diversity and gene flow in *R. norvegicus* in New York.

### Genetic information and invasive rodent management

Informing biosecurity practices based on the genetic information of invasive species is crucial for targeted management actions and the protection of islands [26]. Although such reach was out of the scope of our study, we were able to identify the main dispersal routes of these two invasive rodents of high concern on Cozumel island. Moreover, our findings support there is a continuous introduction of propagules by means of maritime transport and tourism cruises.

One of the most common strategies for the control of invasive species is eradication, which specifically for rodents relies mostly on poisoning. However, this tactic has proven costly and rather inefficient in reducing rodent population in the long term [95]. Aerial or local poisoning (i.e. rodenticide feeding) is not selective, hence extremely dangerous as it can eliminate other species; this threat is more significant in Cozumel as it harbors two endemic rodent species, *Oryzomys couesi cozumelae* and *Reithrodontomys spectabilis* [50,113] and many other endemic vertebrates. Furthermore, given the distribution extent of these invasives on the island and the constant introduction of individuals, poisoning eradication is an unfeasible strategy.

On the other hand, identifying genetic clusters enables defining meaningful spatial units for control management [42]. Indeed, we detected different genetic groups and low connectivity for each species, determined by certain urban and socioeconomic features in accordance with their distinct life history and social organization. Accordingly, we located specific distribution and abundance areas that could be targeted with different management strategies. Constant monitoring and permanent animal trapping should be performed in the main introduction sources, like piers and docks, in urban sites (sanitary landfill, informal commerce) as well as examining the terrestrial and maritime transport vehicles. Very importantly, the strong association of the house mouse with impervious surfaces decreases the chances of a secondary invasion to natural environments, which is vital for systems like Cozumel and the endemic taxa of the island. We should highlight that studies of rodent population ecology and genetics performed in Cozumel over many years have shown that *R. rattus* and *M. musculus* are not found on natural vegetation nor in conserved areas (see [113-115]). Therefore, improving urban planning to regulate future expansions of San Miguel Cozumel is of the outmost importance in order to prevent these invasive species to disperse further on the island. Likewise, it is imperative to make authorities and people aware of the negative effects of these invasives on the native biota and of the associated high potential health risks.

## Acknowledgments

We are grateful with all those that helped during fieldwork, O. Romero-Báez, A. Tobón-Sampedro, N. Rivas, and T. Garrido-Garduño for molecular advice. We deeply thank C. González-Baca, H. Ayuntamiento de Cozumel, Comisión de Agua Potable y Alcantarillado, Comisión Nacional de Áreas Naturales Protegidas, Fundación de Parques y Museos Cozumel, and other institutions, colleagues and inhabitants in Cozumel for their support. GBM acknowledges that this paper was a part of her Bachelor’s thesis in Facultad de Ciencias, Universidad Nacional Autónoma de México.

## Ethical approval

Animal handling followed the procedures allowed by the Mexican Official Norm NOM-062-ZOO-1999 (Technical specifications for the protection, care and use of laboratory animals) [55].

## Supporting information

The online version contains supporting information available as one pdf document, which includes the following:

**Table S1.** Data for *Rattus rattus* and *Mus musculus* individuals sampled on Cozumel Island, México used in this study.

**Table S2.** Microsatellite loci and amplification protocol for both species.

**Table S3.** Ancestral bottleneck results per genetic cluster.

**Figure S1.** Distribution of allele ratio for detection of contemporary bottlenecks.

**Figure S2.** Optimization response curve for *Mus musculus* for the normalized vegetation index (NDVI) variable.

The microsatellite data file is available in the Figshare digital repository and can be accessed at: https:// (will be provided if accepted)

